# Alnustone inhibits Streptococcus pneumoniae virulence by targeting pneumolysin and sortase A

**DOI:** 10.1101/2022.03.07.483358

**Authors:** Can Zhang, Xinyu Wang, Linlin Shi, Baihe Zhan, NaNa Hou, Suohong Liu, Manjie Bao, Gefu Chi, Tianqi Fang

## Abstract

Streptococcus pneumoniae (S. pneumoniae) is a significant Gram-positive opportunistic pathogen responsible for a variety of lethal infections. This bacterium accounts for more deaths from diseases than any other single pathogen worldwide. Distinctively, these symptoms arise despite effective antibiotic therapy. This study unveiled a novel mechanism of resistance to S. pneumoniae infection by targeting pneumolysin (PLY) and sortase A (Srt A), the key virulence factors of S. pneumoniae. Through protein phenotype assays, we found alnustone to be a potent drug that inhibits both PLY and Srt A. Using a PLY-mediated hemolysis assay, we found that albumin can effectively reduce Srt A peptidase activity by blocking PLY oligomerization, thereby directly inhibiting PLY-expressing cytolysis. Co-incubation of S. pneumoniae D39 Srt A with small-molecule inhibitors reduces cell wall-bound Nan A (pneumococcal-anchored surface protein Srt A), inhibits biofilm formation, and significantly reduces biomass. But more interestingly, the protective effect of invasive pneumococcal disease (IPD) on murine streptococcus pneumoniae was further demonstrated. Our study proposes a detailed bacteriostatic mechanism of pneumococcal and highlights the major translational potential of targeting circulating PLY and Srt A to protect against pneumococcal infections. Our results suggest that the antiviral strategy of directly targeting PLY and Srt A with alnustone is a promising treatment option for Streptococcus pneumoniae and that alnustone can be used as an effective inhibitor of PLY and Srt A.

## 1. Introduction

*Streptococcus pneumoniae* (*S. pneumococcus*) is a Gram-positive facultative anaerobic bacterium that causes severe aggressive diseases, including pneumonia, acute otitis media, sepsis, and meningitis, particularly in children and the immunocompromised in the individual ^[1-3]^. Community-acquired bacterial pneumonia is a major diagnosis, treatment disease, with up to 50% of deaths caused by *streptococcus pneumoniae[4]. S. pneumoniae* is also a major cause of community-acquired pneumonia (CAP), with 14.5 million cases and 800,000 deaths annually in children under 5 years of age[5]. However, as antibiotic resistance spreads and expands, treating antibiotics as a routine treatment for *S. pneumoniae* is increasingly being threatened by increased medical costs and mortality[6]. Therefore, new therapeutic strategies to address these challenges are imminent, especially for infections with antibiotic-resistant *S. pneumoniae*. In the pathogenesis of infection, pneumoniae produces a variety of well-characterized virulence factors, including the pore-forming toxins pneumolysin(Ply), Sortase A, capsule (Cps), pneumococcal surface protein PsaA, PspA, PspC, PavA, and PsrP, Neuraminidase and Hyaluronidase (Hyl)[7, 8]. Ply plays an important role in invasive *Streptococcus pneumoniae* infection, and Ply knockout strains have significantly reduced toxic power in mice with nasal and systemic infection models as previous studies have shown. The above results demonstrate that Ply was an important target candidate for drug development to treat *S*.*pneumoniae* infections[9, 10]. Sortase A (Srt A) as another important virulence factor, that was a membrane-anchored transpeptidase expressed by Gram-positive bacteria with Srt A mutant strains able to lead to a greatly reduced ability of infected animals, and Srt A was involved in the colonization and/or pathogenesis of multiple strep species as reported[11, 12]. Therefore, the pressing challenge of microbial infection research is the development of inhibitors and the identification of novel drug targets that can be used for the therapy of infections[13]

Recently, plant extracts have attracted much attention as an alternative strategy against infection. Various plant extracts were used as effective candidates for treating bacterial infections that act on bacterial virulence factors directly to achieve the purpose of reducing bacterial pathogenicity without affecting bacterial evolutionary pressure as in previous studies[14-16]. Alnustone is a non-phenolic diarylheptanoid compound with chemical structures of the type aryl-c7-aryl skeleton isolated from the Alnus pendula (Betulaceae)[17]. Alnustone reportedly exhibits a variety of antiemetic[18], antibacterial[17], anti-inflammatory[19], and anti-hepatotoxic[20]. However, the potential effect of this compound on *S. pneumoniae* infection has not been described. This study revealed that alnustone was a potent Ply and Srt A inhibitor and verified the inhibition mechanism and potential therapeutic effect of the drug on pneumococcal infections through in vivo experiments.

## 2. Materials and Methods

### 2.1. Bacterial strain, cell and culture conditions

*S. pneumoniae* strain D39 serotype 2 (NCTC 7466) was obtained from Dr. David, University of Alabama (Birmingham, U.S.A). Bacteria were grown in Todd-Hewitt broth (THB, HOPEBIO Co, Ltd., Qingdao, China) containing 2% yeast extract (THY) in sealed tubes at 37°C overnight. Bacteria were then harvested and used to inoculate liquid cultures in THB.

The liquid culture was left at 37°C until the OD _600_ reached 0.1 according to the previous culture conditions. Human lung cancer epithelial cells (A549) were purchased from American Type Culture Collection (ATCC, CCL-185, Manassas, VA, USA) and cultured in Dulbecco’s modified Eagle’s medium/high glucose (DMEM; HyClone, Logan, UT, USA). The medium contains 10% FBS(Biological Industries, Kibbutz Beit Haemek, Israel) and 4% penicillin-streptomycin (MRC, Madrid, Spain).

### 2.2. Chemicals

Alnustone (CAS^#^: 33457-62-4, Purity>98%) was commercially obtained from Chengdu Derui Biotechnology Co., Ltd. (Derui, China) and dissolved in dimethyl sulphoxide (DMSO, St. Louis, MO, U.S.A).

### 2.3. Pneumolysin purification

Expression and purification of the pET-28a vector in the experiments for PLY generation. was maintained as described previously[21].

### 2.4. Hemolysis assays

The hemolytic activity assay was slightly modified according to the previous method[22]. Briefly, 10μL of purified Ply protein (1.0 mg/mL) was co-incubated with different concentrations of Alnustone(0, 4, 8, 16, 32, and 64 μg/mL)in 965μL of phosphate-buffer saline (PBS) at 37°C for 30min, and then 25μL of defifibrinated rabbit erythrocyte was added for incubation at 37°C for 8 min until hemolysis was observed. After centrifugation at 12000rpm for 2 min, the value of OD AT 570nm was measured with a microplate reader (Tecan, Melbourne, Austria). PBS was the negative control and double distilled water was a positive control.

### 2.5. Growth curves assay`

*S*.*pneumoniae* strain D39 cultured overnight was expanded in sterile THY at the dilution rate of 1:100 and cultured in an incubator at 37°C until the value of OD at 600nm reached 0.3. The bacteria solution was evenly distributed to a co-culture system containing different concentrations of alnustone (0, 2, 4, 8, 16, and 32μg/mL) respectively. Then, the co-culture solution was cultured at 37°C without shaking, and the OD_600_ value was measured every hour.

### 2.6. Lactate dehydrogenase (LDH) release and live/dead cells assays

A549 cells (type II pneumocytes) were maintained in Dulbecco’s Modified Eagle Medium (DMEM) supplemented with 10% fetal bovine serum. Cultured cells laid into 96-well plates at a density of 3 × 10^4^ maintained in a humidified atmosphere (5% CO_2_ at 37 °C) overnight. Then, various concentrations of alnustone (0, 2, 4, 8, 16, and 32 μg/mL) were added to the cell wells and incubated at 37 °C for 7 hours. DMSO was used as a negative control and triton 0.2% was used as a positive control, respectively. The detached cells were centrifuged at 1000rpm for 10min and 50μL supernatant was mixed with LDH reagent at a ratio of 1:1. After incubation at 37°C for 1h, the absorbance was measured at 450nm with a microplate reader (Tecan, Austria). LDH release levels were analyzed using a Cytotoxicity Detection Kit (Roche, Basel, Switzerland). Purified Ply(1μM)was co-cultured with various concentrations of alnustone at 37°C for 30min, and then A549 cells were incubated into 96-well plates at a density of 3×10^4^well at 37°C for 7h. The centrifugal supernatant (1000rpm,10min) was mixed with 100μL live/dead cells staining reagents in each well and co-cultured at 37°C for 40min according to the instructions of the live/dead cell staining kit (Invitrogen, Carlsbad, CA, USA), then photographed by immunofluorescence microscope(Olympus, Tokyo, Japan).

### 2.7. Immunoblot and oligomerization analysis of Ply

*S. pneumoniae* strain D39 was co-cultured with various concentrations of alnustone (0, 2, 4, 8, 16 and 32μg/ mL) at 37°C. The precipitate after centrifugation (12000 rpm for 5 min) of solution with an absorption value of 1.2 was removed with β-mercaptoethanol (β-ME) in 1×SDS-PAGE loading buffer and heated at 10°C for 8min. Proteins were separated by 12% SDS-PAGE and transferred to a polyvinylidene flfluoride (PVDF) membrane. The membrane was blocked in 5% milk powder (dissolved with TBST), incubated with a rabbit anti-primary antibody (1:1000) overnight at 4°C, followed by incubation with HRP-conjugated secondary anti-mouse antibody (1: 1000, Proteintech) for 2h. Then, targets were detected using ECL reagent (Thermo Scientific, Rockford, IL, USA), and the protein bands were observed using a Tanon-4200 imager (Tanon, Shanghai, China). Purified Ply (dissolved in PBS in a 1:1 ratio) was mixed with various concentrations of alnustone (0, 4, 8, 16, and 32μg/mL) for 40min at 37°C. The cultures with 0.1M alnustone were added with the 5× SDS-PAGE loading buffer without-thiol ethanol at 100°C for 8min. Then the Ply-formed oligomers and monomers were analyzed according to the Previous studies about Western blot analysis.

### 2.8. Sortase activity inhibition assay and biofilm assays

In this study, the inhibition of Alnustone on SrtA peptidase activity was detected by fluorescence resonance energy transfer test (FRET). The supernatant containing Srt A was co-incubated with different concentrations of alnustone at 37° for 30min, and the substrate (Dabcyl-QALPETGEE-Edans) was added, and the fluorescence intensity was detected by microplate analyzer under the conditions of 350nm excitation and 520nm emission after co-incubated at 37° for 1h.

Mid-log-phase bacterial suspension (1 × 10^8^ CFU/mL) was diluted with sterile culture medium THY (THB+2% yeast extract) at a ratio of 1:100, and 10μL bacteria solutions were added to each well of a 24-wells plate with different concentrations of alunstone. The absorption value of D39 culture at 570nm was measured after co-culture without shaking for 10h (0, 2, 4, and 8μg/mL of alunstone) and 20h (0, 4, 8, and 16μg/mL of alunstone) at 37°C under 5% CO_2_ respectively. The plates were stained for 1h for 400μL of 0.1% crystal violet (CV) after washing three times with 1 mL PBS. The OD_570nm_ was measured using a microplate reader, and the bound CV was dissolved in 200μL of 33% glacial acetic acid after the plates were allowed to dry.

### 2.9. Mouse Infection Assays

BALB/c mice (6-8 weeks, female) were purchased from Changsheng Biotechnology Co. Ltd. (Liaoning, China)_and fed with 12h light-dark cycle. All animal experiments were following the guidelines of the ACUC of Jilin University. A mice model of *S. pneumoniae* infection was established by instilling 50μLof PBS containing *S. pneumoniae* strain suspension D39 (2×10^8^CFU) into the left nasal cavity of mice. Mice were injected subcutaneously with 50μL alunstone (25mg/kg body weight) once every 12h for 72h. No treatment was used as a blank control group and the same volume of DMSO was injected as a solvent control group. The survival rate of mice was recorded every 12h within 96h. The BALF was collected and cytokine tested using the ELISA kit instructions (Biolegend, U.S.A) 48 hours after infection. Lung tissue was fixed in a 4% formaldehyde solution for staining with hematoxylin and eosin and for histopathological analysis.

### 2.10. Statistical analysis

Experimental data represent Means ± SEM of at least three independent experiments. Statistically, the Kruskal-Wallis test was used and the difference was statistically significant: *P < 0.05; **P < 0.01; and NS, no significant difference.

## 3. Results

### 3.1. Alunstone inhibits the hemolytic activity of Srt A

Alnustone (Figure 1A) is a yellowish needle crystal, which mainly exists in the flowers of alder in Betulaceae, the seeds of cardamom in Zingeraceae, and the rhizome of yellow root and turmeric. According to the analysis of antibacterial activity test results, alnustone showed a significant bacteriostatic effect on *S*.*pneumoniae* in the range of 4-32μg/ml (Figure 1C), but the co-culture of various concentrations of alnustone (0, 4, 8, 16, and 32μg/ml) with the bacteria did not affect the growth *S. pneumonia* (Figure 1B).

**Figure 1.**
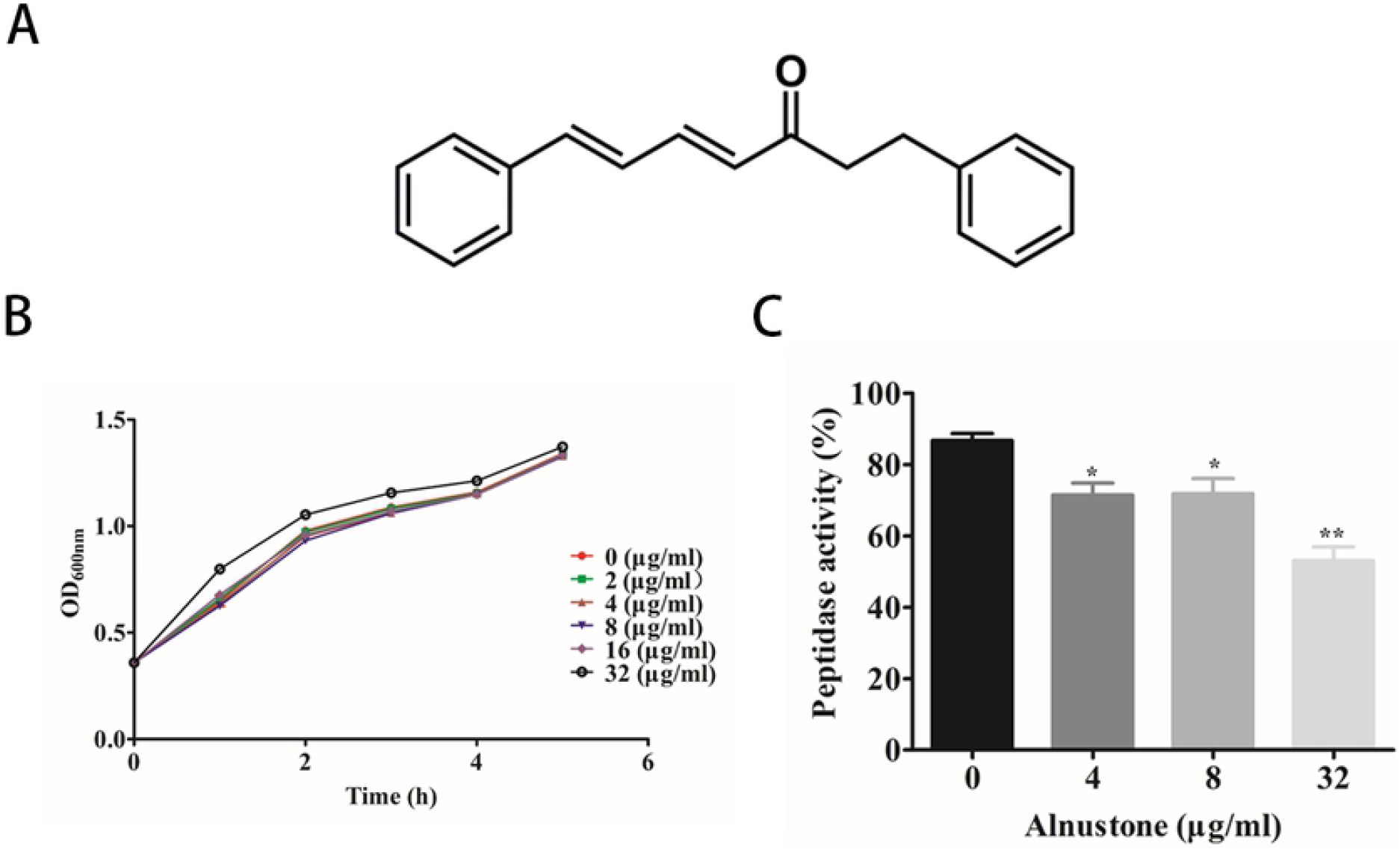
The inhibitory effects of Alnustone on Sortase or the culture supernatant of Streptococcus pneumoniae. (A) Chemical structure of Alnustone. (B) The growth curve of Streptococcus pneumoniae D39 in the presence of different concentrations of Alnustone. (C) The inhibition of SrtADN81 peptidase activity by Alnustone. The supernatant of Streptococcus pneumoniae and various concentrations of Alnustone were incubated for 30 min at 37°C, followed by the addition of SrtADN81 substrates and the fluorescent peptide Dabcyl-QALPETGEE-Edens and incubation for 1 hr. Finally, the fluorescence values of the reaction system were measured. (excitation and emission wavelengths of 350 and 520 nm, respectively). All data are shown as the mean ± SD (n = 3), * indicates P < 0.05, ** indicates P < 0.01. An unpaired two-tailed T-test was used for statistical analysis of the data.

### 3.2. Alnustone the formation of PLY oligomers

Hemolysin (PLY) is one of the important virulence factors of *S. pneumoniae*. PLY is characterized by its ability to form transmembrane macropores in the cholesterol-rich cell membrane, leading to cell lysis and even death. Therefore, we sought to determine whether the small alder molecule could exert its potency by inhibiting the formation of PLY oligomers. Experimental results show that Alnustone inhibited RBC hemolysis in a dose-dependent manner (Figure2 A and B). The expression levels of PLY were the same in the co-culture medium with different concentrations of alnustone and *S. pneumoniae* (Figure 2C). Western blot results showed that protein monomers increased gradually with the increase of alnustone concentration, while oligomers decreased. These results indicated that the formation of PLY oligomers was inhibited with increasing alnustone concentration (Figure 2D). Quantitative density analysis showed that the formation of oligomers gradually decreased with the increase of alder concentration (Figure 2E), which was also consistent with the above results.

**Figure 2.**
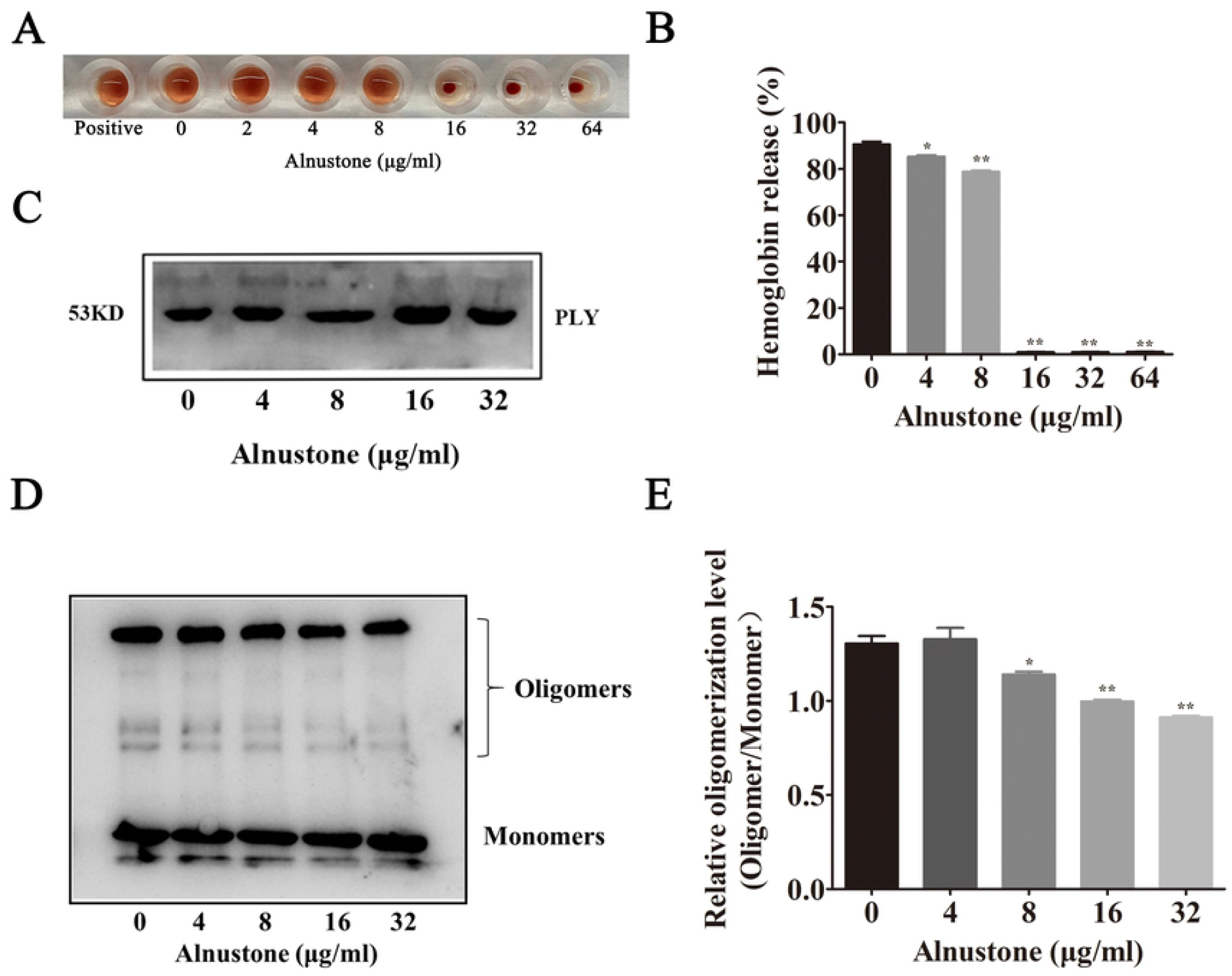
Alnustone inhibits PLY hemolytic activity and the oligomerization of PLY and the localization of NanA. (A) Hemolytic activity of the supernatants from Streptococcus pneumoniae (D39) co-cultured with Alnustone. The sample treated with DD H2O was regarded as a positive control (100% hemolysis). (B) Absorbance value of hemolytic supernatant. (C) Effect of Alnustone on the expression of PLY of bacterial solution precipitation. (D) Inhibitory effect of Alnustone on the oligomerization of PLY. The different concentrations of Alnustone (0, 4, 8, 16, and 32μg/ml) were incubated with purified PLY, and the PLY oligomerization was detected by Western blot analysis. (E)Quantitative analysis of the oligomerization level estimated by ImageJ software (*P < 0.05,**P < 0.001, NS, no significant difference).

### 3.3. Acacetin alleviates PLY-mediated A549 cells injury

As one of the virulence factors of *S. pneumoniae*, hemolysin PLY can damage not only red blood cells but also other cells, such as human alveolar basal epithelial cells (A549) in lung cancer. In this study, the release of lactate dehydrogenase (LDH) was used as an indicator of cytotoxicity. First, it was verified that alnustone alone was not toxic to A549 cells, and different concentrations of the drug did not damage the cells (Figure 3A). However, the release of LDH in the co-culture medium of A549 cells and alnustone decreased with the increase of alnustone concentration (0-16μg/mL) (Figure 3B), which indicated that alnustone had a good protective effect on the cell damage caused by PLY. According to the staining of live dead cells, after co-culture with PLY and A549 cells, under the fluorescence microscope, a large number of cells showed red, indicating a large number of dead cells. Even if there were live green glowing cells, the condition was not very good, but after adding alnustone (16ug/ml), the dead cells were significantly reduced (Figure 3C-3F). In conclusion, alnustone had a significant protective effect on the cell damage induced by PLY cultured in vitro.

**Figure 3.**
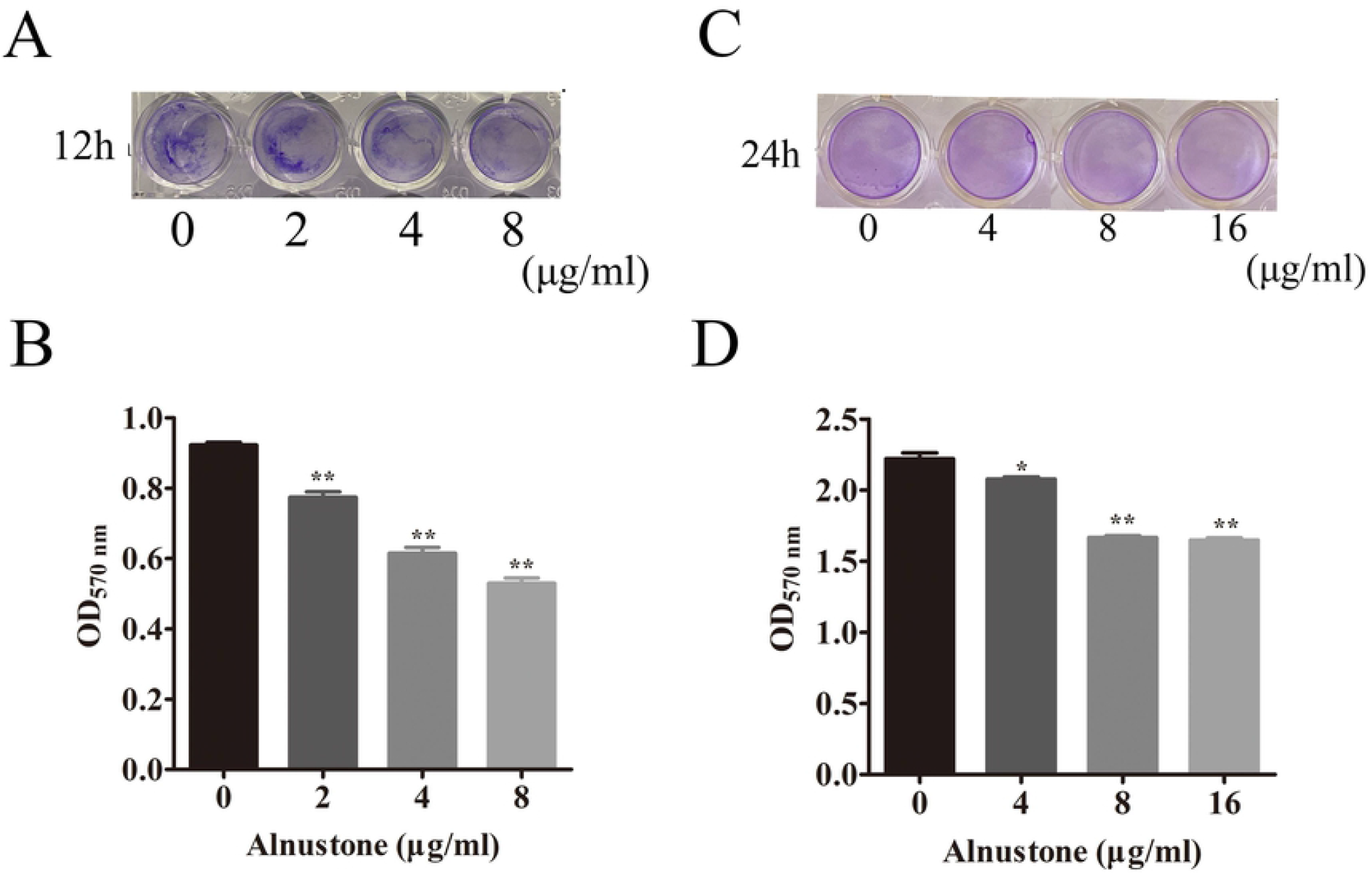
Alnustone inhibits S. pneumococcal biofilm formation and adhesion to human epithelial cells. Streptococcus pneumoniae biofilms were grown in the presence of the indicated concentrations of Alnustone for 12 hrs and 24 hrs. The photograph of a biofilm stained with crystal violet at 12h (A) and 24h (C).Quantification of biofilm formation by measuring bound crystal violet(B, D). The results shown in A are representative of the results from three independent experiments. The bars show the mean values of three independent assays. The error bars indicate S.D. * indicates P < 0.05 and ** indicates P < 0.01 compared with the drug-free group according to 2-tailed Student’s t-tests.

### 3.4. Alnustone impairs pneumococcal biofilm formation and reduces biomass

Small molecule inhibitors can effectively inhibit the formation of *S. pneumoniae* biofilm ^[24]^, which was also proved in this experiment. Biofilm formation also plays an important role in the adhesion of *S. pneumoniae* to host cells. In the crystal violet staining experiment of biofilm, Alnustone showed significant inhibition on the formation of streptococcus pneumoniae biofilm at 12h and 24h respectively (Figure 4A, 4B). The inhibition rate of Srt A peptidase activity was gradient-dependent. Biofilm formation also plays an important role in the adhesion of *S. pneumoniae* to host cells.

**Figure 4.**
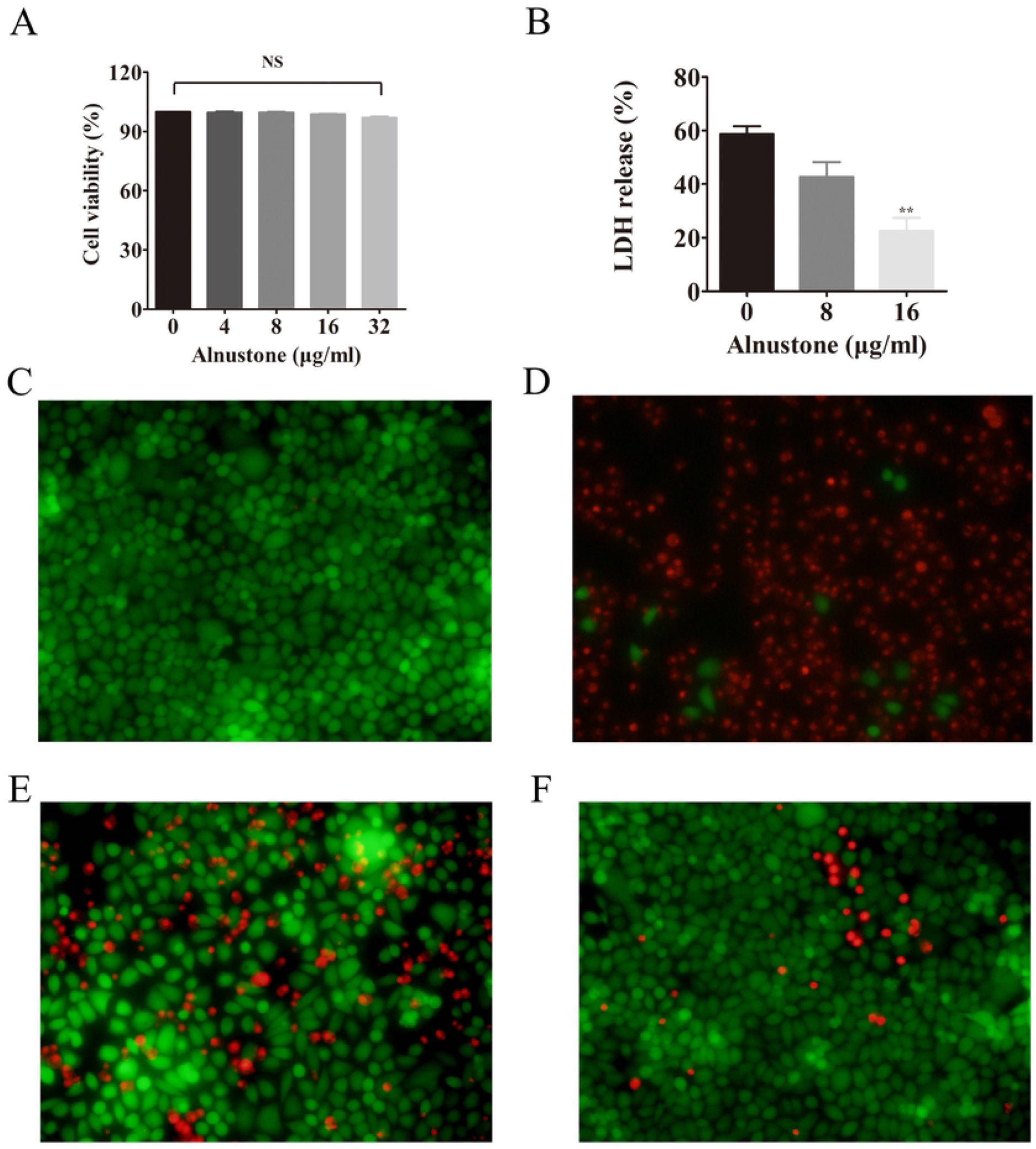
Alnustone alleviates PLY-mediated A549 cells injury. (A) The cytotoxicity of Alnustone on A549 cells. (B) LDH was released in the coculture system of A549 cells and purified PLY with various concentrations of Alnustone. Live/dead A549 cells assay by immunofluorescence microscopy after incubation with PLY in the presence of acacetin. (C) Uninfected A549 cells. (D) A549 cells were incubated with PLY. (E) A549 cells were incubated with PLY in the presence of 8 μg/ml. (F) A549 cells were incubated with PLY in the presence of 16 μg/ml. The results shown in C, D, E, and F are representative of the results from three independent experiments. The bars show the mean values of three independent assays. The error bars indicate S.D. * indicates P < 0.05 and ** indicates P < 0.01 compared with the drug-free group according to 2-tailed Student’s t-tests.

### 3.5. Acacetin attenuated Streptococcus pneumoniae virulence in mice

In the mice *pneumoniae* model was established by *S*.*pneumoniae* nasal attack method, and the protective effect of alnustone for mice was observed by treatment with various alnustone. Consistent with previous studies, the infected mice showed significant pathological changes 48 hours after infection compared with the uninfected mice. The macroscopic lesion observation showed that the lung injury of mice treated with different concentrations of alnustone was significantly reduced (Figure 5A). According to the HE staining sections of lung tissues, compared with the untreated group, alnustone has a good protective effect on pneumonia in mice, as shown in the figure 5A: After treatment with alnustone, the lung inflammation of mice has significantly reduced These results were consistent with the pulmonary ocular changes in the front, thus indicating that alnustone can effectively protect mice from s. pneumoniae infection in vivo.

**Figure5.**
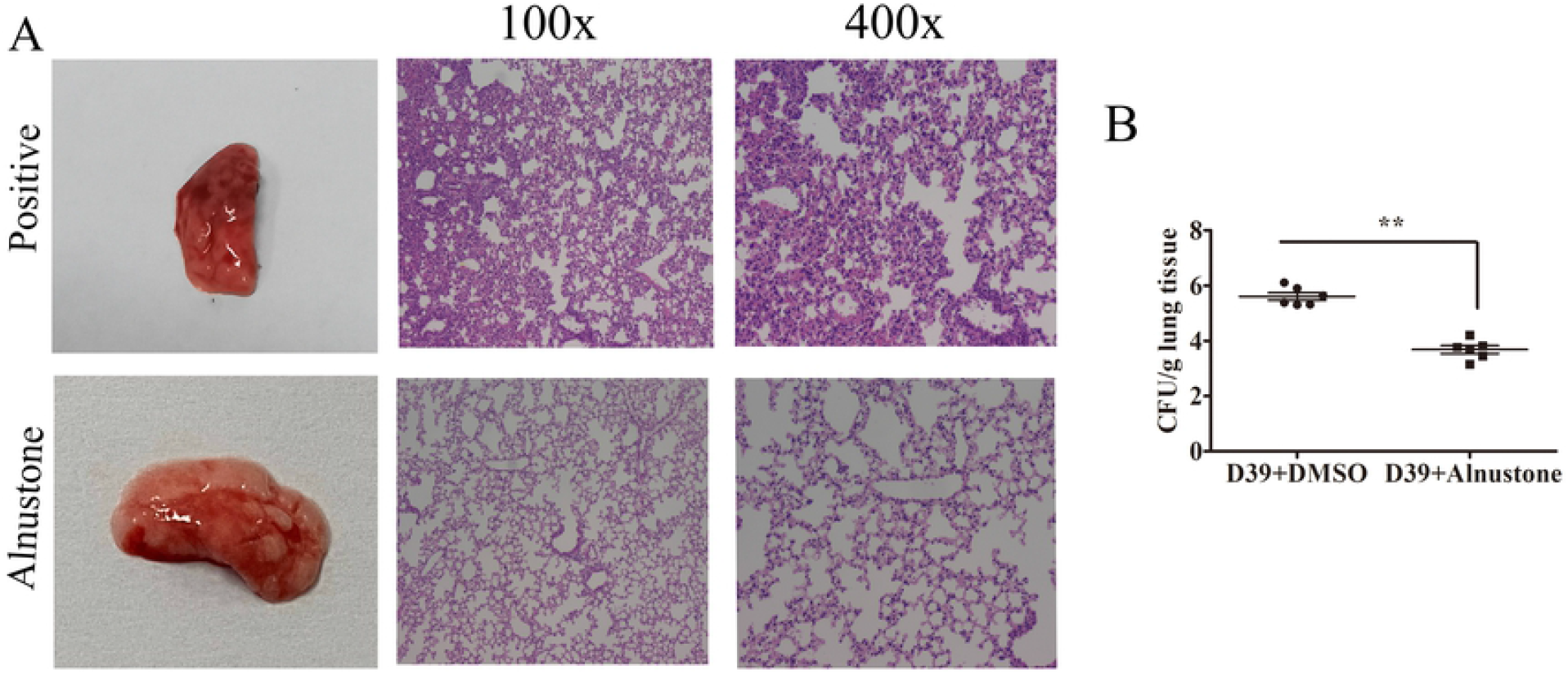
Alnustone protects mice against S. pneumoniae. BALB/c (n=6) were inoculated with S.pneumoniae via the intranasal route and treated subcutaneously with PBS or alnustone. The gross pathological changes and histopathology(A) of lung tissue after 48 hours post-inoculation were assessed. The results shown in (A) are representative of the results from three independent experiments. The bars in (B) show the mean values of three independent assays. The error bars indicate S.D. * indicates P < 0.05 and ** indicates P < 0.01 compared with the drug-free group according to 2-tailed Student’s t-tests.

## Discussion

Recently, the treatment of *S. pneumoniae* infection meets more serious challenges with the emergence and widespread of various antibiotic resistance in recent years[23]. Currently, alternative strategies for bacterial virulence factors without affecting their viability have gradually been concerned by researchers. It has demonstrated that small molecule inhibitors can significantly weaken pneumococcal virulence in vitro and in vivo[9]. The cytolytic function of Ply was related to blood flow, microbial colonization, immune escape, and host cell apoptosis in pneumococcal pneumonia[24]. In addition, another important virulence factor of *S. pneumoniae* was associated with the anchoring of surface proteins in Gram-positive bacteria. It also has been demonstrated that inactivation of the Srt A gene significantly attenuated surface protein display and bacterial virulence[25].

In this study, we selected the important virulence factor pneumolysin (PLY) and Srt A as the targets of anti-infective for drug screening. Hemolysis assays were used to screen for natural compounds with the ability to antagonize Ply, and it is determined that their directly diminished effect on the pathogenicity of pathogenic bacteria. The hemolysis results showed that Alnustone at lower concentrations (4 μg/mL) could directly inhibit Ply-mediated hemolysis activity, while not inhibiting bacterial growth. The effect of Alnustone on the expression level of PLY expression levels and oligomerization in vitro were further analyzed by immunoblotting. The results showed that alderone suppresses the oligomerization process of PLY in a dose-dependent manner without affecting the normal hemolysin expression level, which should be a direct mechanism of action for antagonizing the hemolysin activity. In addition, we found that Alnustone was also a potent inhibitor of Srt A. Srt A was essential for bacterial colonization and biofilm formation via the LPETG motif according to previous studies. It has been shown that pneumococcal bacteria in biofilms trend towards increased antimicrobial drug resistance due to lower antibiotic penetration into biofilm structures partly. At the same time, the protective effect of alnustone against *S. pneumoniae* infection was confirmed by the Srt A peptidase activity inhibition assay and the biofilm formation inhibition assay. The results of this study indicate that the forming ability of biofilm and adhesion ability to epithelial cells was diminished significantly in the case of the coculture with *S. pneumoniae* D39 and alnustone. The protection of the drug in vivo was evaluated through a mice intranasal infection with *S. pneumoniae*. The results showed that alnustone can significantly reduce the pathological damage and the wet/dry proportion of the lung while inhibiting autoinflammatory responses and improving the survival rate of infected mice. These findings further demonstrate that alnustone has dual roles in vivo and in vitro to provide effective protection against *S. pneumoniae* infection.

In conclusion, Alnustone was found to neutralize the pore-forming activity of Ply effectively and to inhibit oligomers forming Ply. It can reduce the damage of Ply to alveolar epithelial cells (A549) significantly. In addition, alnustone can gradient-dependent reduce the peptidase activity of SrtA, but also significantly affect the adhesion of shoulder support bacteria to host cells, while inhibiting the formation of biofilm. Alnustone as a double target inhibitor of Ply and Srt A can reduce the damage caused by bacteria hemolysin to the body, which lays a good preliminary test foundation and provides lead compounds for the development of the new drug as double target drug-resistant pathogenic bacteria infection based on inhibiting bacterial pathogenicity and resistance.

## Acknowledgments

This work was supported by the National Natural Science Foundation of China (grant NO. 81861138046,32102723 and 31972742)

## Author contributions

Gefu Chi and Tianqi Fang designed the study; Can Zhang and Xinyu Wang performed the experiments; Linlin Shi, Baihe Zhan, NaNa Hou, Suohong Liu, and Manjie Bao contributed reagents/materials/analysis tools; Can Zhang wrote the paper; Tianqi Fang reviewed the manuscript.

